# “Rapid SARS-CoV-2 Detection Using High-Sensitivity Thickness Shear Mode Sensors”

**DOI:** 10.1101/2024.05.22.594713

**Authors:** Sahera Saleh, Habib Alkalamouni, Karen Antar, Pierre Karam, Jit Muthuswamy, Hassan Zaraket, Massoud L Khraiche

**Affiliations:** Department of Experimental Pathology, Immunology & Microbiology, Faculty of Medicine, American University of Beirut, Beirut, Lebanon; Center for Infectious Disease Research, Faculty of Medicine, American University of Beirut, Beirut, Lebanon; Neural Engineering and Nanobiosensors Group, Biomedical Engineering Program, Maroun Semaan Faculty of Engineering and Architecture, American University of Beirut, Beirut 1107 2020, Lebanon; Chemistry Department, American University of Beirut, Beirut, Lebanon; Neural Microsystems Laboratory, School of Biological and Health Systems Engineering, Arizona State University (ASU), Tempe, AZ, United States

## Abstract

The COVID-19 pandemic, caused by the SARS-CoV-2 virus, has emphasized the urgent need for accurate and readily available diagnostic tools. Conventional diagnostic methods, such as reverse transcription real-time polymerase chain reaction (RT-qPCR), are often labor-intensive and time-consuming, which highlights the necessity for rapid point-of-care diagnostic solutions. This study introduces an innovative, low-cost, and highly sensitive diagnostic platform for swift COVID-19 detection. Our platform utilizes the mass sensing properties of thickness shear mode (TSM) transducers to detect and quantify the SARS-CoV-2 nucleocapsid protein through polyethylene glycol (PEG)-based chemistry (1). To confirm surface functionalization and evaluate the effects of the virus lysis buffer, we employed surface characterization techniques including Digital Holographic Microscopy (DHM), Scanning Electron Microscopy (SEM) with Energy-Dispersive X-ray spectroscopy (EDX), and Raman spectroscopy. Sensitivity tests with heat-inactivated SARS-CoV-2 samples demonstrated a sensitivity of about 0.256 Hz/TCID50/mL and a limit of detection (LOD) of roughly 150 TCID50/mL. Specificity was verified through cross-reactivity testing. Our detailed characterization and sensitivity analysis underscore the platform’s reliability, making it a promising candidate for efficient and accessible COVID-19 diagnosis at the point of care.

## 1 Introduction

The coronavirus infectious disease, known as COVID-19, is caused by the severe acute respiratory syndrome coronavirus 2 (SARS-CoV-2) which emerged in late 2019 in Wuhan, China (2, 3, 4). It rapidly evolved into a global pandemic, marking the third major spillover event of a coronavirus from animals to humans in the past two decades. Unlike its predecessors, COVID-19 is highly transmissible, with various transmission modes (5, 6, 7, 8, 9). Despite considerable advancements in vaccination efforts and public health measures, the impact of COVID-19, particularly with the emergence of new variants, continues to be a significant public health concern and a socioeconomic burden (10, 11). Accurate and fast testing remains a crucial tool for effectively managing COVID-19 and mitigating its impact (10, 12, 13).

Currently, two primary types of COVID-19 diagnostic tests are available: viral antigen detection via immunodiagnostic techniques and nucleic acid amplification method (14, 15). The latter uses nucleic acid amplification techniques (NAAT) like RT-qPCR and can also employ antigen testing to detect viral proteins. However, the sensitivity of rapid antigen tests is lower than RT-qPCR, making them less effective for COVID-19 diagnosis (16). Additionally, rapid antigen tests do not provide quantification of the viral load. Whereas, RT-qPCR have its own limitations, including the time-consuming nature of the process and the requirement for specialized equipment and trained personnel (17). This is where the necessity for rapid, quantitative diagnostic point of care tests becomes crucial. Rapid diagnostic tests offer a faster and more accessible alternative to qRT-PCR, enabling quicker identification of SARS-CoV-2 infections (18).

In this work, we present a quantiative point-of-care sensing platform for rapid COVID-19 detection. The scalability of the proposed sensing approach and related instrumentation lends itself for future high-throughput detection and deployment. Our approach utilizes a thickness shear mode (TSM) transducer for mass sensing. Surface acoustic wave (SAW) devices, such as those based on thickness shear mode (TSM) resonators, have emerged as powerful tools for biosensing applications. TSM devices operate by propagating acoustic waves along the surface of a piezoelectric crystal, and their sensitivity to mass changes makes them highly suitable for detecting biomolecular interactions (5, 6, 7, 19). The TSM biosensors offer a distinct advantage due to their label-free detection capability, allowing near real-time monitoring of molecular binding events without the need for additional chemical labels. This inherent simplicity not only streamlines experimental procedures but also minimizes potential interference with the binding kinetics. Additionally, TSM sensors exhibit high sensitivity and precision, enabling the detection of minute changes in mass or viscoelastic properties on the sensor surface. The compatibility of TSM biosensors with a wide range of biological samples, from small molecules to larger biomolecules, further extends their versatility in biosensing applications (20), (21). The proposed platform utilizes polyethylene glycol (PEG)-based chemistry for the detection and quantification of the SARS-CoV-2 nucleocapsid antigen protein, offering several advantages including improved sensitivity and specificity. (1, 22). The surface chemistry employed in this platform has been characterized using multiple techniques to ensure its reliability and efficacy in capturing the target analyte. Through this work, we aim to address the pressing need for efficient and accessible diagnostic tools for COVID-19. By presenting a novel mass sensing approach coupled with a custom surface chemistry targeted towards the viral protein of interest, this study contributes to the ongoing efforts in combating the pandemic and addresses the need for rapid point-of-care testing to mitigate the impact of the pandemic.

## 2 Materials and Methods

### 2.1 Surface Chemistry

Gold TSM sensors (resonating frequency of 10 MHz) were purchased from openTSM Q-1 (Novaetech, Italy). Prior to functionalization, the chips underwent a cleaning process using a strong acid piranha solution (H2O2 /H2SO4 in a 3:1 v/v ratio). The cleaning process involves immersing the chips in the solution for a duration of 5 minutes, effectively eliminating organic material deposited on the surface, then rinsing with deionized (DI) water and air drying for 15 minutes (23). The sensors were then treated with a 1M sodium hydroxide (NaOH) solution for 20 minutes. For functionalization and immobilization of antibodies for capturing the SARS-CoV-2 nucleocapsid protein, Thiol-PEG-carboxyl was employed. The thiol group enables chemisorption on gold surfaces. Thiol-PEG-carboxymethyl coatings can be customized to achieve specific surface density. A 1 mg/mL solution of thiol-PEG-carboxyl (HS-PEG-COOH, MW 5K) (Laysan Bio Inc., Alabama, USA) is introduced to the surface of the sensor and incubated in DI water for 2 hours at room temperature. Multiple DI water washes were performed to eliminate free PEG chains. The COOH group of COOH-PEG-SH was activated through a two-step reaction using 0.4 mg 1-Ethyl-3-(3-dimethylaminopropyl)carbodiimide (EDC), 0.6 mg N-Hydroxysuccinimide (NHS), 0.1 M 2-(N-morpholino) ethane sulfonic acid (MES), and 0.5 M sodium chloride (NaCl) for an incubation time of 14 minutes at a pH ranging from 5.5 to 6.0. EDC is then quenched by adding 1.4 µL of 20 mM beta-mercaptoethanol. Multiple washes with a phosphate buffer solution (PBS) (0.1M sodium phosphate, 0.15 M NaCl, adjusted to pH 7.2-7.5) were then performed (24).

Following functionalization and activation, 20 µL of SARS-CoV-2 nucleocapsid protein (NP) (E8R1L) fragmented mouse monoclonal antibody f-mAb (Cell Signaling Technology Inc., Massachusetts, USA) are introduced to the sensor surface after diluting in PBS and incubated for 2 hours at room temperature. Unreactive groups were then quenched by adding excessive ethanolamine (ETA), ending the amine conjugation reaction followed by multiple PBS washes Figure 1.

**Figure 1:**
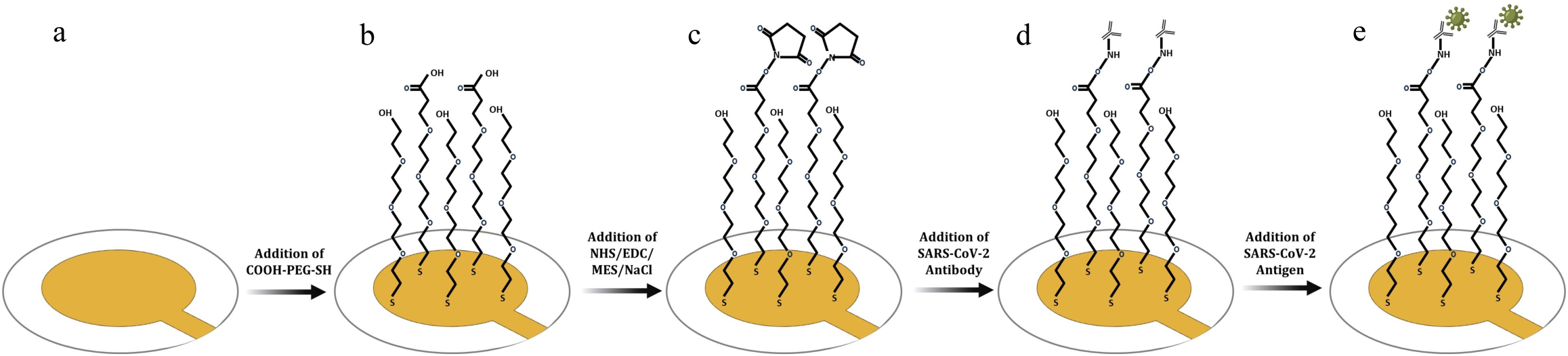
Sequential steps involved in the surface functionalization. (a) Sensor is cleaned with distilled water (DIW) after piranha and sodium hydroxide treatment. (b) Addition of 1 mg/mL solution of thiol-PEG-carboxyl (HS-PEG-COOH, MW 5K). (c) Activation of COOH-PEG-SH and addition of beta-mercaptoethanol to quench 1-Ethyl-3-[3-dimethylaminopropyl] carbodiimide (d) Addition of SARS-CoV-2 Nucleocapsid Protein (E8R1L) Mouse monoclonal antibody mAb and incubation for 2 hours at room temperature. (e) Addition of SARS-CoV-2 heat inactivated virus sample to the surface.

### 2.2 Virus Lysis

A series of characterization techniques were employed to determine the lysis buffer that preserves the surface chemistry of the coated sensor. Choosing the appropriate buffer is contingent upon numerous factors, including viral characteristics, pH, detergents, chelating agents, buffer components, and enzymes. We investigated two lysis buffers formulations. Lysis buffer #1 was prepared according to the formulation described by Raziq et al., 2021 (25), consisting of 27.5 mM Tris-HCL, a buffering agent to maintain pH stability, 12.5 mM Ethylene diaminetetraacetic acid (EDTA), as a chelating agent to sequester divalent cations, stabilizing viral components during lysis, 1.5% Triton, a nonionic detergent that ruptures the viral envelope liberating the viral components and 0.1% Sodium Sodium Dodecyl Sulfate (SDS), an anionic detergent for protein disintegration. Lysis buffer #2 was prepared according to the formulation described by Singh et al., 2022 (26), consisting of 50 mM NaH2PO4H2O, a buffering reagent to maintain pH stability, 300 mM NaCl, added to establish an isotonic environment, ensuring virus particle integrity and 10 mM Imidazole, a buffering reagent to maintain pH conditions. The inactivated virus sample is mixed with the selected lysis buffer (1:3 v/v ratio) and incubated at 56°C for 30 minutes to achieve effective lysis (27). This formulation emphasizes buffering reagents for pH stability, establishing an isotonic environment with NaCl to preserve virus particle integrity, and using Imidazole as an additional buffering reagent.

### 2.3 Surface Characterization

#### 2.3.1 Scanning Electron Microscopy (SEM) with Energy-Dispersive X-ray spectroscopy (EDX)

Surface characterization techniques were employed to gain insight into the stability of the functionalized surface of the gold quartz sensor. In this work, we characterized the surface using Scanning Electron Microscopy (SEM) with Energy-Dispersive X-ray spectroscopy (EDX). SEM images were obtained using an accelerating voltage of 5 kV after sputter-coating with gold. EDX analysis confirmed the integrity of the functionalized surface.

#### 2.3.2 Reflection Digital Holographic Microscope

Reflection Digital Holographic Microscope (DHM®, LyncéeTec, Switzerland), that employs holographic interferometry technology to capture a live holographic image, was employed to generate a real-time 3D reconstruction of the surface and to quantify its roughness after different steps of functionalization (23, 28). Monitoring the surface roughness allows for observing and assessing the binding events on the surface of the sensor and evaluating the surface topography.

#### 2.3.3 Raman spectroscopy

Raman measurements were carried out in the backscattering configuration using a 532 nm laser line from an Ar-ion laser. Incident light was focused to a spot size of around 1 μm using a confocal microscope with an x100 objective. The laser power on the sample surface was only 3 mW for all measurements to avoid contributions from laser heating. The scattered light was analyzed using a triple monochromator and a Peltier-cooled charge-coupled detector (CCD). A 50-μm slit was used to reduce the spectral broadening of the spectrometer to below 0.5 cm^2.

### 2.4 TSM Assay Sensitivity

The sensitivity of the developed immunosensor was evaluated by comparing its frequency response to different concentrations of SARS-CoV-2 lysates (Isolate USA-WA1/2020, heat inactivated) and bovine serum albumin (BSA) in PBS (negative control). To test the device’s overall sensitivity, incremental BSA concentrations were added on a bare gold quartz sensor initially wetted by PBS. To test its sensitivity to SARS-CoV-2, the lysed heat-inactivated viral sample of a known concentration was added over a fully functionalized sensor, and subsequent dilutions were performed. The shift in frequency corresponding to each concentration was then measured over a 5-min period after the signal stabilizes for both tests. This sensitivity test provided insights into the immunosensor’s performance in a diagnostic context. Furthermore, we quantified the limit of detection (LOD), defined as the minimum concentration of analyte at which a consistently observable positive test line is obtained. The World Health Organization (WHO) has established “acceptable” LOD values of approximately 1000 TCID50/ml, but 100 TCID50/mL as “desirable” target for SARS-CoV-2 *in vitro* diagnostics (4).

The relationship between the sensor’s response and analyte concentration was established by generating a calibration curve. This curve allowed us to determine the analyte concentration in an unknown sample by comparing its sensor response to the calibration curve. To ensure the study’s reliability and validity, a vehicle control using PBS was performed in parallel.

The sensitivity of the immunosensor was determined by analyzing the slope of the calibration curves for both BSA and SARS-CoV-2 concentrations. These sensitivity values provide valuable information about the immunosensor’s performance and its ability to detect specific proteins at different concentrations.

### 2.5 Real-Time Reverse Transcription PCR (RT-qPCR)

To determine the Ct values corresponding to various dilutions of SARS-CoV-2, we performed a RT-qPCR using the Reliance One-Step kit from BioRad. The reaction utilized primers designed for the RNA-dependent RNA polymerase (forward primer: CTCACCTTATGGGTTGGGATTA, reverse primer: GTTTGCGAGCAAGAACAAGTG, probe: FAM-TGATAGAGCCATGCCTAACATGCT-BHQ1) Cycling conditions adhered to the manufacturer’s protocol.

## 3 Results and Discussion

### 3.1 Comparative Analysis of Effect of Lysis Buffers on Integrity of Surface Chemistry Integrity using Surface Topography

Digital Holographic microscope (DHM) images of the gold quartz sensor were acquired at different stages of the functionalization process and the corresponding roughness values were measured. Figure 2 presents DHM images of a bare sensor showing an initial average surface roughness of 2.22 nm after cleaning. Upon introducing 1 mg/mL COOH-PEG-SH and subsequent washing with DI water, we observed a notable increase in surface roughness with the average roughness value rising to approximately 4.84 nm. This elevation in average roughness can be attributed to the molecular adsorption and attachment of COOH-PEG-SH molecules to the sensor’s surface, modifying its topographical features. The subsequent addition of NP mAb further increased the roughness to approximately 48.18 nm, indicating possible successful immobilization of SARS-CoV-2 N mAb on the functionalized sensor surface. To investigate the impact of different lysis buffers on the adhered SARS-CoV-2 N mAb, buffer #1 and buffer #2 were applied to the surface-bound antibodies, followed by thorough washing and drying. Surface roughness measurements revealed that the application of buffer #1 resulted in an average roughness of 50.28 nm, while buffer #2 yielded a slightly lower average roughness of 36.64 nm. This variation may be attributed to differences in the interaction of the lysis buffers with the immobilized antibodies.

**Figure 2:**
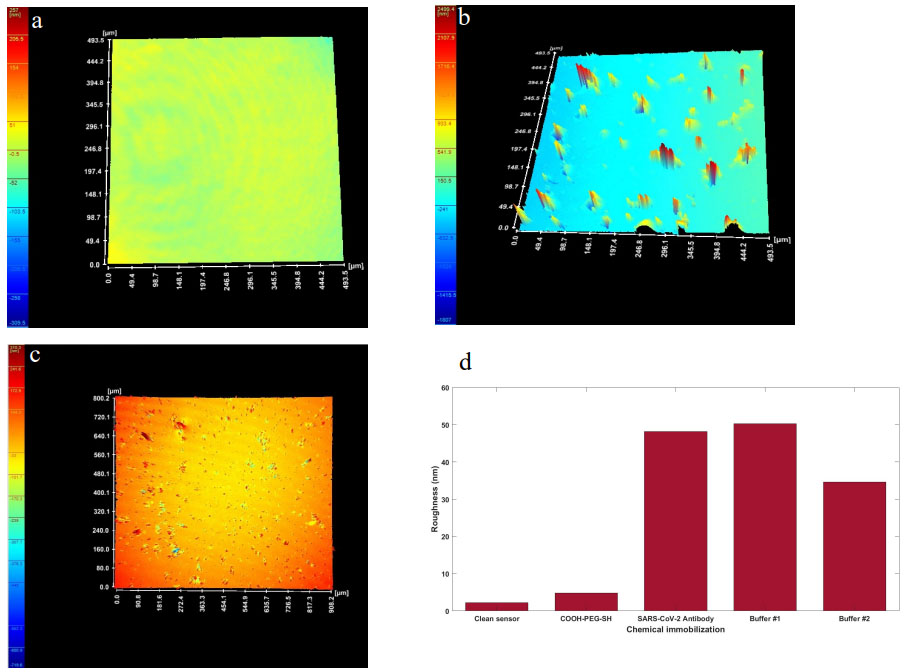
Digital holographic microscope image of (a) bare gold surface (b) sensor surface after SARS-CoV-2 nucleocapsid protein (E8R1L) mouse monoclonal antibody mAb immoblization (c) sensor surface after BSA coating. (d) Changes in surface roughness as measured by digital holographic microscopy after different stages of the chemical immobilization process.

### SEM and EDX Analysis

SEM images and EDX spectroscopy were also utilized to examine potential disruptions to the functionalized surface upon the addition of buffer #1 and buffer #2 on top of the SARS-CoV-2 NP mAb. As shown in Figure S1 (a), EDX analysis of the gold quartz sensor after introducing COOH-PEG-SH identified elements such as Silicon (Si), Gold (Au), Oxygen (O), Carbon (C), and Sulfur (S), confirming the integrity of the functionalized surface. Si appeared as we are working on a quartz sensor, and Au is from the gold electrode surface of the quartz crystal. Carbon, Oxygen, and Sulfur are indications of the presence of thiol-PEG-carboxymethyl. After adding SARS-CoV-2 NP mAb and buffer #1, EDX analysis showed the presence of nitrogen (N), confirming the stable amide bond formation between the antibody and COOH-PEG-SH in Figure S2. Following the addition of SARS-CoV-2 NP mAb and buffer #2, we identified nitrogen (N), sodium (Na), and chloride (Cl), providing evidence of the antibody’s presence and the stable amide bond between the antibody and the COOH-PEG-SH following activation through the EDC/NHS reaction in Figure S3. These results confirm successful binding of SARS-CoV-2 N mAb and demonstrate that neither buffer #1 nor buffer #2 disrupted the functionalized surface.

### Raman Spectroscopy

We investigated the effect of buffer formulation on the integrity of the surface chemistry. Both buffers provide needed buffering capacity, which is essential for maintaining pH stability during the lysis process. The choice, among other factors, depends on the sensitivity of the Thiol-PEG-carboxyl coating to ionic strength, potential interactions with metal ions, and the impact of detergent type on the surface. Lysis Buffer #1, the use of Triton as a nonionic detergent might be milder on the surface compared to anionic detergents, potentially minimizing disruption of the Thiol-PEG-carboxyl coating. On the other hand, for Lysis Buffer #2, NaCl, establishing an isotonic environment, may help preserve the structural integrity of the Thiol-PEG-carboxyl coating during the lysis process. Compatibility testing was done to possible adverse affect of buffer formulations on the Thiol-PEG-carboxyl coating and other surface functionalities. Figure 3 illustrates the Raman spectroscopic analysis of three sensors representing different stages of our immobilization – (1) sensor with COOH-PEG-SH activation and SARS-CoV-2 NP (f-mAb) binding, (2) sensor with SARS-CoV-2 N mAb binding and Buffer #1 application and (3) sensor with SARS-CoV-2 N mAb binding and Buffer #2 Application. The three sensors exhibited similar peak locations, particularly with an overlap observed in the first two peaks within the Raman shift range of 200 to 1400 rel.cm⁻¹. This overlap suggests that the initial functionalization of the sensor surface with COOH-PEG-SH, activation using EDC/NHS, and the subsequent attachment of NP mAb (f-mAb) did not significantly alter the Raman-active functional groups on the surface. However, a notable difference was observed in the Raman shift range of 1000 to 2000 rel.cm⁻¹. In this range, f-mAb and Buffer #1 displayed a similar curve shape for the amide bond at 1653 rel.cm⁻¹. Conversely, Buffer #2 exhibited a distinct curve shape at 1653 rel.cm⁻¹. Also, the intensity ratio from Buffer 1 increases slightly as the wavenumber increases, while the intensity ratio from Buffer 2 decreases as the wavenumber increases. In principle, a Raman signal extending to high wavenumbers/frequencies indicates the presence of light element bonds such as hydrogen and carbon, this indicates Buffer 1 contains these bonds, while Buffer 2 does not. The Raman spectroscopy data suggests that Buffer #1 maintains the the integrity of our functionalized surface (COOH-PEG-SH) and the attachment of the NP mAb to the surface.

**Figure 3:**
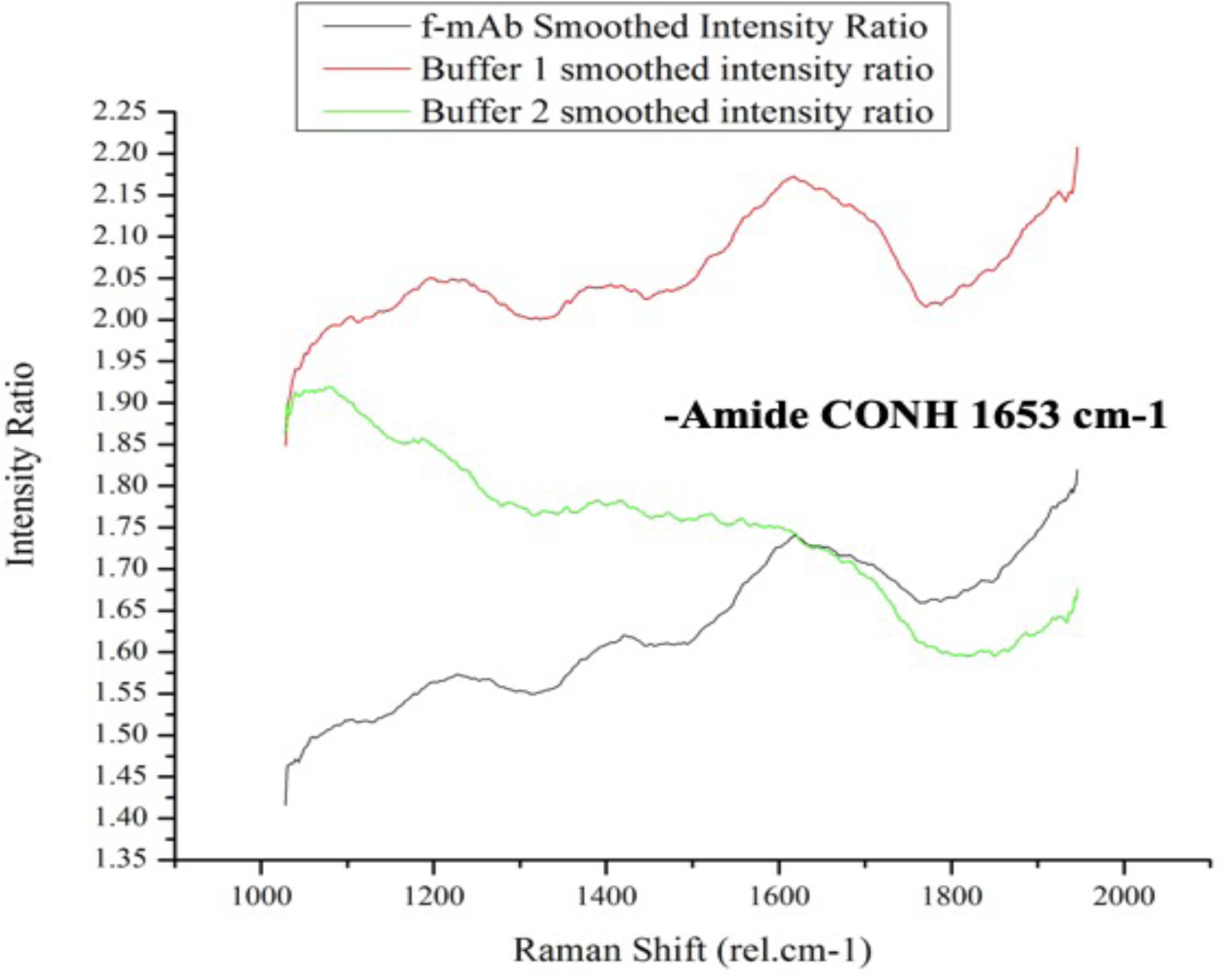
Raman spectroscopy results: Black curve is of sensor having COOH-PEG-SH activated using EDC/NHS and immobilized with SARS-CoV-2 NP mAb. Red curve is of a dried sensor after introduction of buffer #1 on top of SARS-CoV-2 NP mAb bonded to activated COOH-PEG-SH using EDC/NHS. Green curve is of a dried sensor after introduction of buffer #2 on top of SARS-CoV-2 N mAb bonded to activated COOH-PEG-SH using EDC/NHS.

### 3.2 Assay Sensitivity and Limit of Detection

To evaluate the device’s sensitivity to the different viral loads of the lysed heat-inactivated SARS-CoV-2, the frequency response was measured for a series of dilutions of the viral concentrations ranging from 5.33 x10^4^ TCID50/ml down to 5.33 TCID50/ml (Figure 4, Table 1). qRT-PCR was also done in parallel for each viral concentration to determine the corresponding Ct value. To compute the device sensitivity, a linear regression analysis was performed on the logarithmic values of concentration (in TCID50/mL) and the logarithmic values of the average change in frequency (Hz). The slope of the obtained linear regression curve was found to be approximately 0.513. This indicates that for each logarithmic unit increase in concentration (e.g., a 10-fold increase in concentration in TCID50/ml units), the average increase in frequency is approximately 0.513 logarithmic units or 0.03164 Hz/TCID50/mL. The device sensitivity observed in our analysis is of significance in clinical settings. A sensitivity of 0.513 suggests that the TSM COVID-19 sensor is capable of accurately detecting and quantifying SARS-CoV-2 over a wide range of viral loads. This feature is crucial for precise diagnosis and monitoring of viral infections.

**Figure 4:**
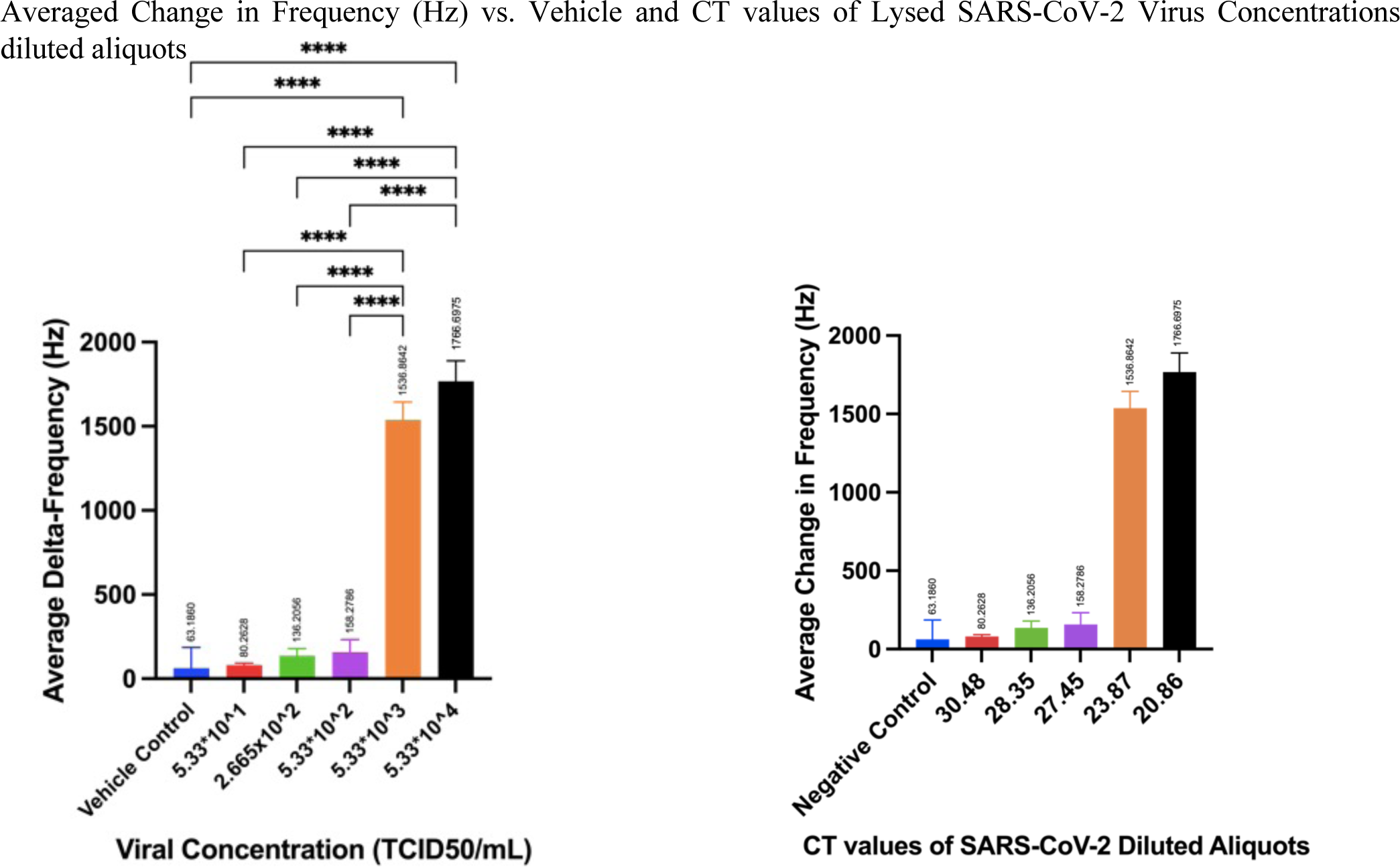
Average change in frequency and the corresponding standard deviation for the vehicle control (DIW and PBS), as well as different viral concentations (left) and the corresponding Ct (right, measured via RT-PCR) value for lysed heat-inactivated SARS-CoV-2 virus samples.

**Table 1:**
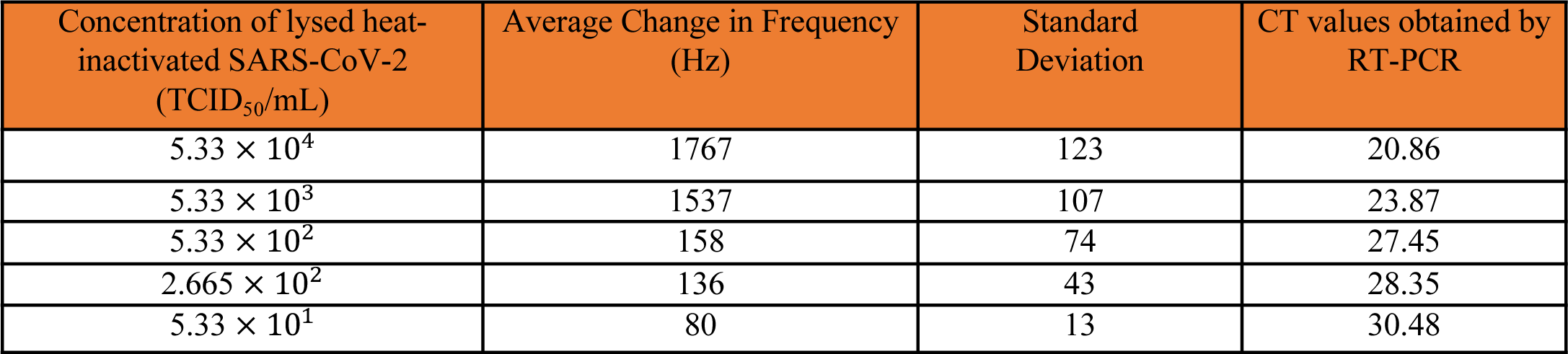
Average change in frequency (Hz) as a function of concentration of lysed heat-inactivated SARS-CoV-2, and CT values obtained by RT-PCR test.

The Limit of Detection (LOD) is a critical parameter for a sensor or assay, indicating the lowest concentration of the target analyte that can be reliably detected above the background. Given the sensitivity calculated earlier, which is approximately 0.03164 Hz/TCID50/mL, and assuming a standard deviation of the blank of 50 Hz (as mentioned in a previous response), we calculate the LOD be 149.36 TCID50/mL. Our sensor demonstrated a linear response when correlating the logarithm of the average change in frequencies with varying concentrations of inactivated virus Figure 5. This observed relationship suggests the potential for accurate quantification of the viral load in tested samples, providing valuable insights into the extent of viral presence and shedding in clinical specimens.

**Figure 5:**
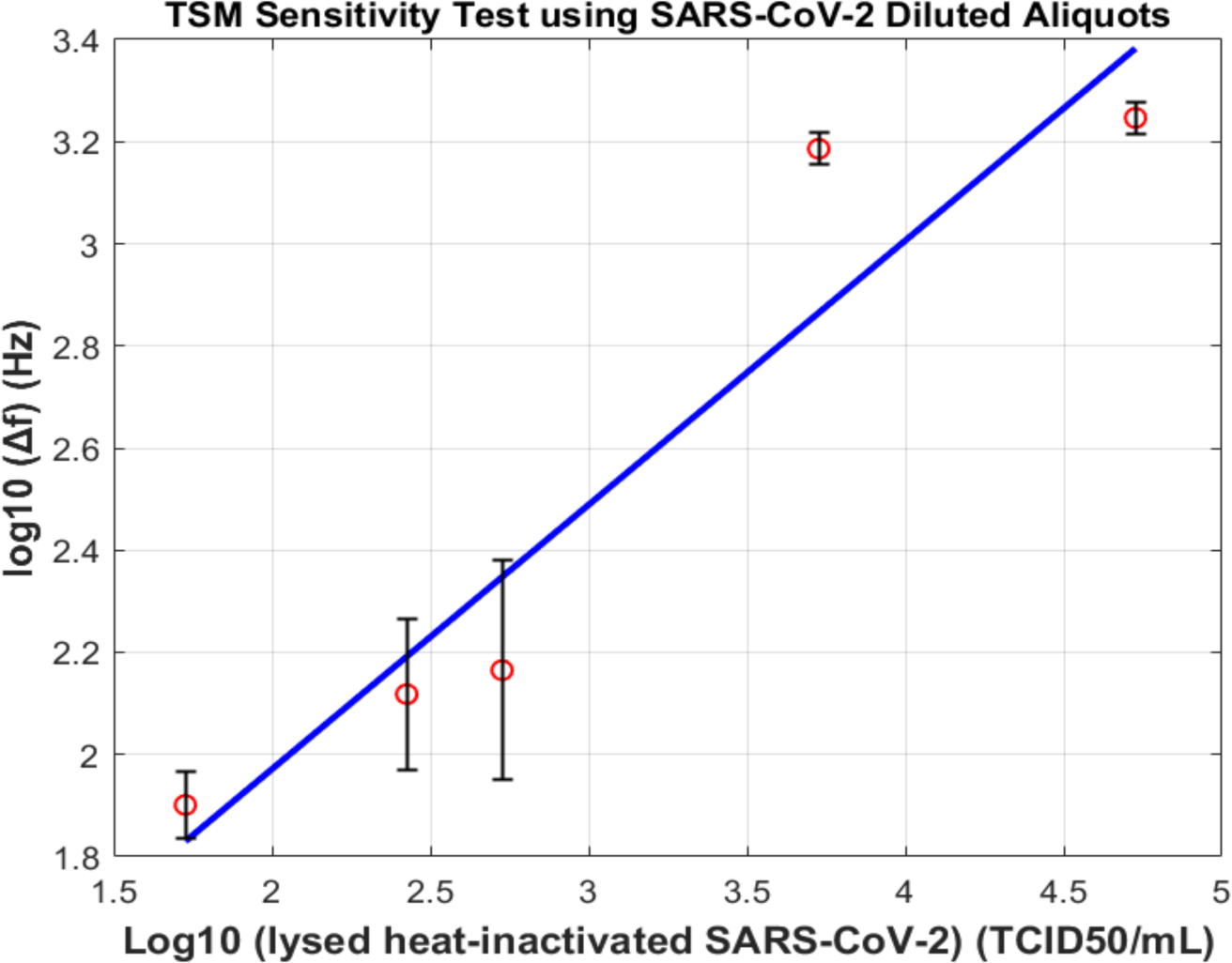
Linear regression of logarithmic values of average Δf (Hz) of all replicates vs. logarithmic values of SARS-CoV-2 inactivated virus concentrations (TCID50/ml). R^2^= 0.9053.

### 3.3 Cross-reactivity

Influenza A virus is a common respiratory virus, like SARS-CoV-2, and can cause similar symptoms and co-circulate during the same period. Testing the biosensor against influenza A virus helps assess its specificity in a clinical setting where multiple respiratory viruses may coexist. For this test, a heat-inactivated Influenza type-A viral sample with an initial concentration of 4.37 × 10^7 TCID50/mL. The Influenza A virus was then lysed using a lysis buffer as previously described. The cross-reactivity test followed the surface functionalization steps with the substitution of Influenza A lysate at the end instead of the SARS-CoV-2 lysate. The specificity of the sensor was examined by observing its binding behavior towards Influenza A antigen. The results of the cross-reactivity test, as depicted in Figure 6, demonstrated that the change in frequency after washing away the Influenza antigen was notably higher than the frequency change after washing the excess SARS-CoV-2 antibody from the surface. While TSM primarily measures frequency changes, it also provides information about energy dissipation. Changes in dissipation can influence the frequency response. Binding should drop the frequency, but the introduction of Influenza A virus lysate can induce changes in the viscoelastic properties of the surface or the surrounding medium, it might lead to an increase in dissipation and, consequently, an increase in frequency. On average, the frequency increased by 80.23 (St Dev =30Hz). The absence of cross-reactivity in the cross-reactivity test is a significant finding. It underscores the specificity of the developed sensor for SARS-CoV-2 and its ability to accurately differentiate the target virus from unrelated or similar pathogens. This specificity is of paramount importance in diagnostic applications, as it helps prevent false-positive results. Moreover, cross-reactivity testing contributes to the accuracy of epidemiological data related to COVID-19. Reliable data on the prevalence and spread of SARS-CoV-2 are crucial for effective public health interventions, resource allocation, and decision-making. By minimizing false-positive results, cross-reactivity testing ensures that individuals are accurately diagnosed and appropriately managed. It is though good to note that while Influenza type A serves as a reasonable choice for cross-reactivity testing, there are considerations associated with using it in this context as it represents only one type of respiratory virus. The cross-reactivity with Influenza type A may not necessarily predict the biosensor’s behavior against a broader spectrum of respiratory viruses. The results might not be fully representative of the diversity of viruses encountered in clinical practice.

**Figure 6:**
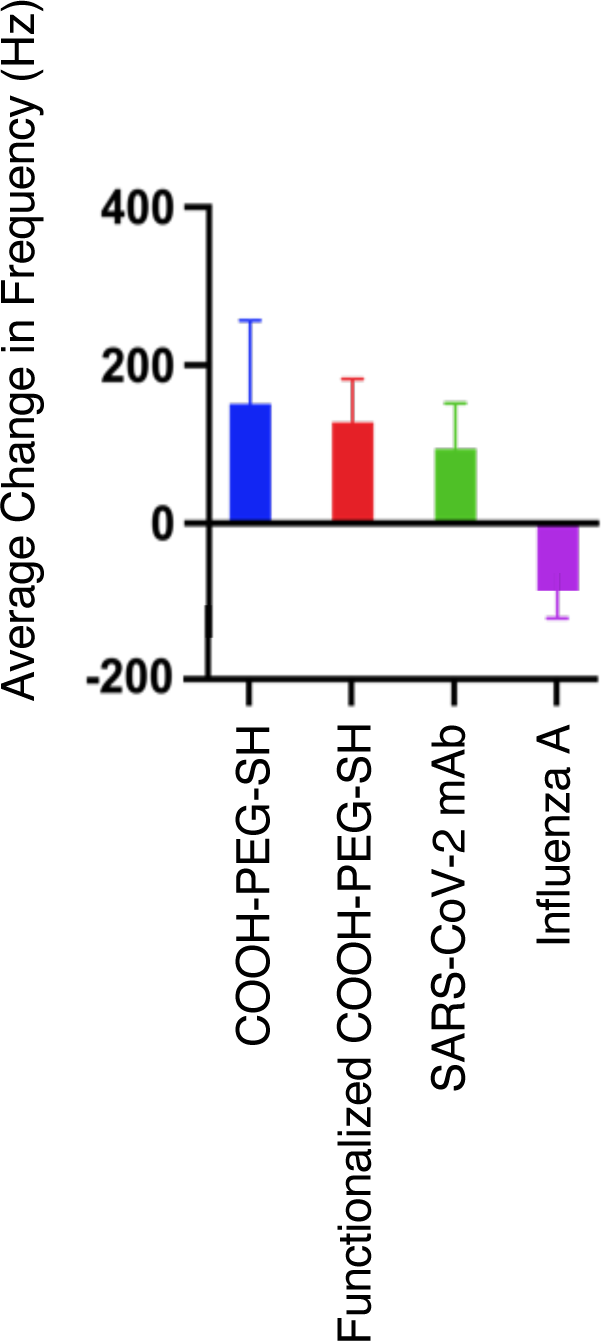
Average Δf (Hz) upon introduction of 2.97 x 10^7^ TCID50/mL lysed Influenza A inactivated virus for cross-reactivity testing. The change in frequency after washing the influenza virus lysate away from the sensor is negative.

## 4 Conclusion and Future Directions

Acoustic technologies have shown great promise in both imaging and sensing applications, enhancing diagnostic capabilities (29, 30). This research introduces a highly promising method for the rapid and specific detection of the SARS-CoV-2 virus, through the development of a rapid, quantitative immunosensor. Our approach stands out as a novel and effective means for diagnosing COVID-19, offering significant advancements in sensitivity and specificity. The study’s strength lies in its comprehensive surface characterization, confirming the efficacy of the functionalization process and validating the reliability and robustness of the sensor surface. Notably, our immunosensor demonstrates a calculated sensitivity of approximately 0.256 Hz/TCID50/mL, enabling accurate detection and quantification of SARS-CoV-2 across a broad range of viral loads. Moreover, the establishment of the Limit of Detection (LOD) at 149.36 TCID50/mL provides a critical benchmark for the diagnostic capabilities of our platform.

While our results are promising, it is essential to acknowledge several limitations and potential challenges associated with the proposed sensor platform. One limitation is the reliance on heat-inactivated viral samples for sensitivity testing, which may not fully represent the complexity of clinical samples. Future research should prioritize validation using clinical samples to ensure the sensor’s performance under real-world conditions. Additionally, the cross-reactivity testing conducted with Influenza type A represents only one type of respiratory virus. Further studies with a broader spectrum of respiratory viruses are needed to fully assess the sensor’s specificity. Addressing these limitations requires ongoing research efforts. Strategies for future research include refining the sensor design to enhance its performance with clinical samples, conducting comprehensive validation studies in diverse populations, and expanding cross-reactivity testing to include a wider range of respiratory pathogens. Moreover, efforts to optimize assay protocols, improve sensor stability, and address potential interference factors will be crucial for advancing the practical utility of the sensor platform. Moving forward, our platform holds promise not only for COVID-19 diagnosis but also for addressing diagnostic needs beyond the pandemic. By providing a cost-effective and user-friendly solution, our platform has the potential to revolutionize healthcare infrastructures, particularly in resource-limited settings and community-based testing initiatives. Despite the challenges ahead, continued research and innovation in sensor technology offer hope for more efficient and accessible diagnostic solutions, ultimately contributing to improved public health outcomes.

In summary, our research significantly contributes to the development of a specific and efficient diagnostic tool for the rapid detection of SARS-CoV-2. As we navigate through the challenges and limitations, our commitment to advancing sensor technology remains steadfast. Through collaborative efforts and continued innovation, we can harness the full potential of our platform to address current and future healthcare challenges, making a meaningful impact on global health security.

## Chemistry and Surface Characterization - Supp Data

**Figure S1:**
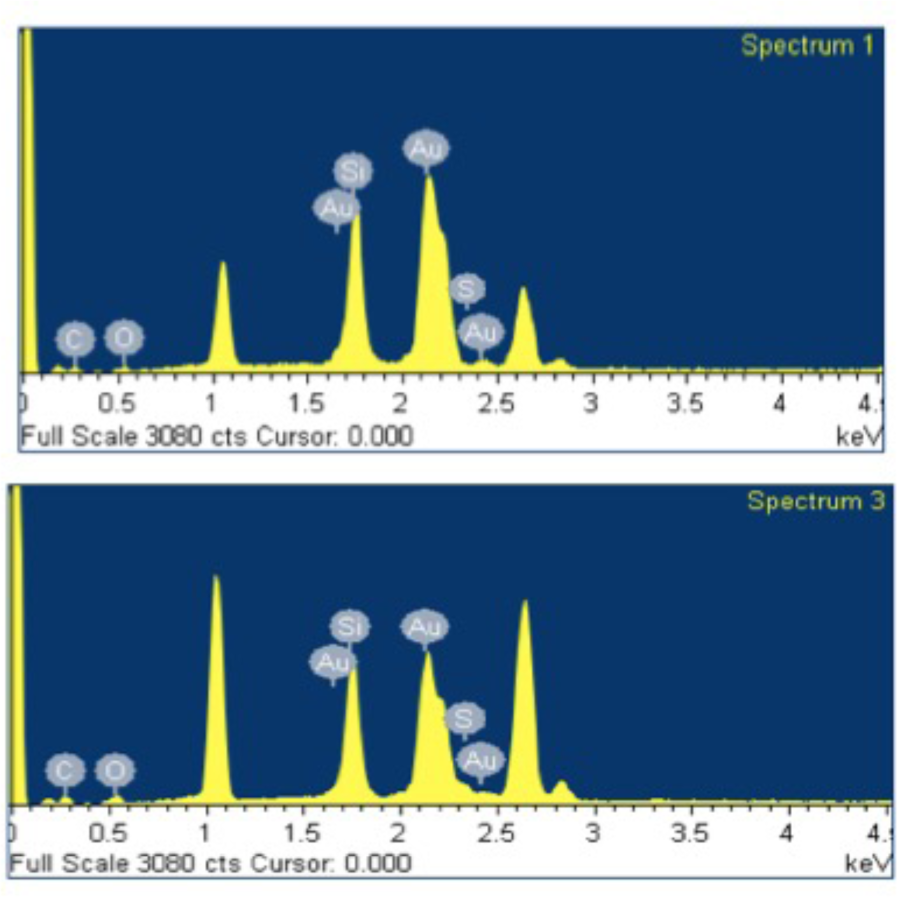
EDX analysis of a washed and dried sensor having a layer of COOH-PEG-SH.

**Figure S2:**
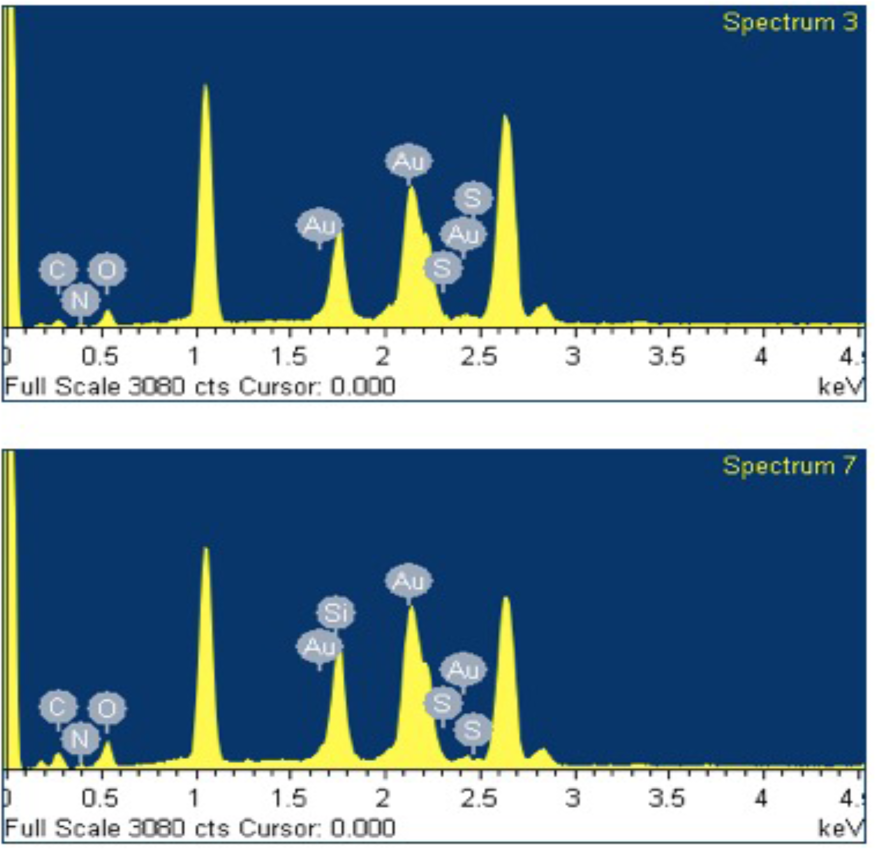
EDX analysis of a washed and dried sensor having buffer #1 added on top of immobilized mAb over crystals of activated functionalized COOH-PEG-SH.

**Figure S3:**
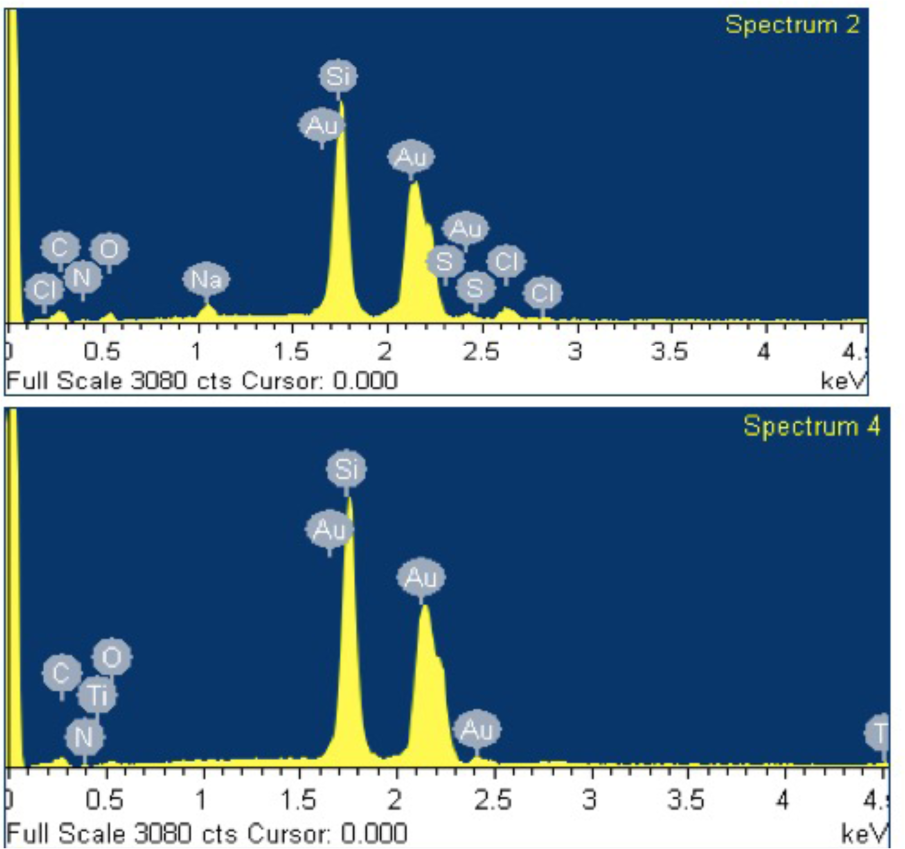
SEM and EDX analysis of a washed and dried sensor having Buffer #2 added on top of immobilized mAb over crystals of activated functionalized COOH-PEG-SH.

